# Single-animal, single-tube RNA extraction for comparison of relative transcript levels via qRT-PCR in the tardigrade *Hypsibius exemplaris*

**DOI:** 10.1101/2024.03.15.585302

**Authors:** Molly J. Kirk, Chaoming Xu, Jonathan Paules, Joel H. Rothman

## Abstract

The tardigrade *Hypsibius exemplaris* is an emerging model organism renowned for its ability to survive environmental extremes. To explore the molecular mechanisms and genetic basis of such extremotolerance, many studies rely on RNA-sequencing (RNA-seq), which can be performed on populations ranging from large cohorts to individual animals. Reverse Transcription Polymerase Chain Reaction (RT-PCR) and RNA interference (RNAi) are subsequently used to confirm RNA-seq findings and assess the genetic requirements for candidate genes, respectively. Such studies require an efficient, accurate, and affordable method for RNA extraction and measurement of relative transcript levels by quantitative RT-PCR (qRT-PCR). This work presents an efficient single-tardigrade, single-tube RNA extraction method (STST) that not only reliably isolates RNA from individual tardigrades but also reduces the required time and cost for each extraction. This RNA extraction method yields quantities of cDNA that can be used to amplify and detect multiple transcripts by quantitative PCR (qRT-PCR). The method is validated by analyzing dynamic changes in the expression of genes encoding two heat-shock-regulated proteins, Heat-Shock Protein 70 β2 (HSP70 β2) and Heat-Shock Protein 90α (HSP90α), making it possible to assess their relative expression levels in heat-exposed individuals using qRT-PCR. STST effectively complements existing bulk and single tardigrade RNA extraction methods, permitting rapid and affordable examination of individual tardigrade transcriptional levels by qRT-PCR.

**SUMMARY:** This work presents a rapid RNA extraction and transcript level comparison method for analyzing gene expression in the tardigrade *Hypsibius exemplaris.* Using physical lysis, this high-throughput method requires a single tardigrade as the starting material and results in robust production of cDNA for quantitative Reverse Transcription Polymerase Chain Reaction (qRT-PCR).

## INTRODUCTION

Tardigrades are small multicellular animals renowned for their ability to survive extreme conditions that are lethal to most other forms of life^1^. For example, these animals can survive nearly 1000-times the dose of ionizing radiation that is lethal to humans^2–10^, nearly complete desiccation^11–15^, freezing in the absence of added cryoprotectants,^16–18^ and, in their desiccated state, even the vacuum of space^19, 20^. Owing to their unique capacity for survival in extreme environments, these animals have become foundational models for understanding extremotolerance in complex, multicellular organisms^1, 21–23^.

Stable genetic manipulation of these remarkable animals, including transgenesis and germline gene modification, has remained elusive until recently^24, 25^. As such, most experiments to reveal molecular mechanisms of extremotolerance are performed through transcriptional profiling via RNA sequencing. Many valuable and informative RNA sequencing data sets exist for tardigrades under various extreme conditions, ranging from radiation^8, 9, 26–28^, heat stress^29^, freezing stress ^12, 17, 29^, and desiccation^27, 30–33^. Some of these studies have utilized bulk RNA extraction and purification methods to illuminate our molecular understanding of extremotolerance. However, bulk extraction of RNA transcripts from many animals prevents analysis of variation in gene expression between individuals, thus missing the potential richness of more refined data sets. Importantly, these studies often analyze heterogeneous populations of animals that include both animals that survive environmental stressors and those that do not. As such, these studies are confounded by averaging expression data from multiple and potentially dramatically different response states. To address this issue, Arakawa et al., 2016^34^ developed an elegant low-input RNA-seq pipeline that applies an RNA extraction kit followed by a linear PCR amplification step using single^34–36^ or multiple^26–28, 30, 37, 38^ animals as input. These studies have been foundational to our understanding of tardigrade extremotolerance^22^. Interestingly, this protocol has also been applied to qRT-PCR using seven animals as starting material^39^.

In most model organisms, having identified potential targets via RNA-seq, qRT-PCR is then performed to confirm transcriptional changes identified by RNA-seq and assess the expression time course of candidate genes in a high-resolution manner. To test the function of identified genes, such studies are often followed by RNAi-mediated knockdown of molecular targets^40, 41^ and analysis of extremotolerant capacity^12, 42^. The efficacy of each RNAi knockdown is typically confirmed by qRT-PCR by directly monitoring the decrease in transcript abundance. However, RNAi is a labor-intensive process in tardigrades as each dsRNA must be delivered via manual microinjection of individuals^40, 41^. Owing to the low throughput nature of this strategy, a rapid, low-cost RNA extraction method adapted for qRT-PCR from single animals would be highly valuable for tardigrade research. Although previous methods have been developed to extract RNA from single tardigrades, these protocols have not combined their extraction with qRT-PCR, instead relying on optical density-based methods^12, 41, 42^. Motivated by these challenges, we sought to develop a protocol that reliably yields RNA in quantity and quality that can be used for qRT-PCR from single *H. exemplaris*.

Adapted from a single-animal RNA extraction protocol developed for *Caenorhabditis elegans*^20^, STST is optimized for *H. exemplaris*. The extraction method consists of six rapid freeze-thaw steps, physically disrupting the cuticle, allowing RNA extraction and subsequent cDNA synthesis. The STST method decreases extraction time by more than 24-fold compared to bulk RNA extraction methods, as described by Boothby, 2018^43^, and by 30% compared to single tardigrade RNA extraction kits, as described by Arakawa et al., 2016^44^. Further, the number of sample-experimenter interactions is decreased from 5 to only 1 compared to RNA extraction kit preparations, thus reducing the risk of contamination by exogenous ribonucleases. When querying for highly expressed genes, the STST method produces sufficient cDNA for 25 quantitative RT-PCR reactions per single tardigrade, requiring only 1 μL of the total 25 μL cDNA volume per reaction. However, template concentrations need to be empirically determined for lower abundance transcripts.

We evaluated the efficacy of the STST method for analyzing dynamic changes in gene expression by investigating the differential expression of the genes encoding heat-shock protein-90α (HSP90α) and heat-shock protein 70β2 (HSP70β2) in response to short-term heat-shock at 35°C for 20 minutes. Both HSP70β2 and HSP90α in most eukaryotic organisms are rapidly upregulated following short-term heat-shock exposure (20 minutes)^45^. Analysis in *H. exemplaris* revealed that both the HSP70β2 and HSP90α-encoding RNAs extracted from single heat-treated tardigrades showed statistically significant increases in expression following short-term heat exposure. These findings demonstrate that the STST protocol can be used to analyze dynamic changes in gene expression in individual animals over time.

The STST extraction method should complement existing experimental methods such as RNA-seq by facilitating rapid and inexpensive RNA extraction and subsequent comparison of transcript levels by qRT-PCR. This method will also be valuable for assessing the efficiency and penetrance of RNAi in manually injected individuals more quantitatively than optical density alone. Finally, owing to their similar cuticular structures and physical characteristics, it is likely that this method will also be effective for analyzing gene expression in other tardigrade species^46^.

### PROTOCOL

For detailed tardigrade and algal culturing procedures, refer to McNuff et al. 2018^47–49^.

1. Sterilization of Spring Water

1.1. Pour 2 L of spring water from a 5-gallon water jug (see Materials Section for Specifics) into a 2 L autoclave-safe glass bottle.
1.2. Place the cap on the autoclave-safe bottle and seal with a small amount of autoclave tape. Do not tighten the bottle; just place the cap on top.
1.3. Autoclave the spring water for 50 minutes on a wet cycle with no drying step.
1.4. Allow the water to come to room temperature and seal the cap firmly before storing it at room temperature.
2. Glass micropipette pulling (with a pipette puller)

2.1. Secure a glass micropipette (O.D. 1 mm, I.D. 0.58 mm, Length 10 cm) on a micropipette puller. Avoid contact with the heating filament, as this will alter the pipette shape and damage the filament. The pulling of the pipette will need to be determined empirically for each filament and pipette puller. However, to serve as a starting point for optimization, use 78°C and a single pull step of pull weight of 182.2 grams.
2.2. Allow the filament to heat and gravity to separate the glass micropipette into two glass micropipettes with sharp points (**Figure 1b**).
2.3. Store these pulled glass micropipettes in a closed 100 mm petri dish with wax or clay to hold them in place and prevent the sharp tips from breaking.
3. Glass micropipette pulling (without a pipette puller)

3.1. Light a Bunsen burner or other controlled flame source on a low setting.
3.2. Take a glass micropipette with one end in each hand.
3.3. Hold the center of the glass micropipette over the flame until the glass begins to melt. Then, rapidly pull the two ends apart. This will create two very delicate sharp tips.
3.4. Lightly break the tip with a pair of sterile fine forceps.
3.5. Store these pulled glass micropipettes in a closed 100 mm petri dish with wax or clay to hold them in place and prevent the sharp tips from breaking.
4. RNA extraction

4.1. Obtain 0.5 L of liquid nitrogen in a cryo-safe container. *CAUTION: Liquid nitrogen is cryogenic and may cause burns if exposed to skin or eyes. When handling, use protective clothing, splash goggles, nitrile gloves, cryo-gloves, a lab coat, and closed-toed shoes. Ascertain that the container is liquid nitrogen safe before transporting the liquid. Using an ethanol-dry ice bath for this step may also be possible*.
4.2. Make cDNA synthesis master mix: a 10 µL solution containing 1 µL of random hexamer primer, 2 µL of DNase, 4 μL of 5x RT Buffer, 1 µL Enzyme Mix, 1 µL of H2O, and 1 μL of 10 mM dNTPs. Store this solution on ice.
4.3. Prepare Tardigrade Lysis Buffer (5 mM Tris (pH=8), 0.5% (v/v) Detergent 1, 0.5% (v/v) Detergent 2, 0.25mM EDTA in sterile nuclease-free Water). This solution can be stored on the bench top for 6 months. However, maintain sterility and avoid potential RNAse-contaminating sources.
4.4. Aliquot enough lysis buffer for extractions (2 μL/ tardigrade).
4.5. Add RNAse inhibitor to the Tardigrade Lysis Buffer solution to a final concentration of 4 units/μL (U/μL).
4.6. Vortex and spin down the solution at room temperature on a bench-top centrifuge at a speed of 2000 xg for 5 seconds before storing the solution on ice.
4.7. Remove as many tardigrades as needed for your experiment from a culture using a sterile filter-tipped P1000 pipette and place them in a sterile 35-mm Petri dish. Any number of tardigrades may be processed in this way. Usually, three tardigrades per condition are processed for extraction.
4.8. Wash the tardigrades three times, using 1 mL of autoclaved sterile spring water and a sterile filter-tipped P1000 pipette. Slowly pipetting them up and down helps to remove algal contaminants.
4.9. Using a dissecting microscope at 25x to 50x magnification, transfer a single tardigrade from this washed culture to a new sterile 35-mm petri dish using a sterile filter-tipped P10 pipette.
4.10. Use a sterile filter-tipped P200 pipette to wash the single tardigrade in 100 μL of sterile nuclease-free water. This wash step is used to further remove contaminants, including ribonucleases.
4.11. Transfer the washed tardigrade to the bottom of a clean, sterile PCR tube in 1-2 μl of sterile nuclease-free water using a sterile filter-tipped P10 pipette, carefully ensuring the tardigrade is not stuck to the side of the tip.
4.12. Visualize the tardigrade under a dissecting microscope at 25x magnification.
4.13. To facilitate water removal of water, break the tip of the pulled glass micropipette lightly outside of the tube. The bore should be big enough to pull up the water but not the tardigrade.
4.14. Using the capillary action of a pulled glass micropipette, remove water until the animal is surrounded by a small bubble of water approximately two tardigrade lengths in diameter.
4.15. Monitor the water removal process via the dissecting scope to ensure the water level is appropriate and the tardigrade remains hydrated. **Figure 1c** offers an example of how much water to remove. *NOTE: This is a critical step. A small bubble of water will surround the tardigrade to prevent it from drying out, but as much excess water as possible should be removed to prevent dilution of the lysis buffer. For an example of the remaining water levels, please refer to* ***Figure 1c***.
4.16. Immediately after removing the water, add 2 μL of Tardigrade Lysis Buffer to the bottom of the tube, briefly vortex, and centrifuge the tube at room temperature for 5 seconds at 2000 x g on a tabletop centrifuge.
4.17. Immediately place the samples containing the tardigrades into a PCR tube rack and ensure that they are held tight by the rack.
4.18. Grip the rack using a pair of long coarse forceps and gently dip the rack containing the samples into the liquid nitrogen until fully frozen. (**Figure 1d**).
4.19. Remove the rack from the liquid nitrogen and immediately place it on ice. Allow the sample to thaw (this should take 45 seconds to 1 minute total). Monitor the sample every 15 seconds by removing it from the ice and visibly inspecting it. Once the sample is visibly transparent, move on to the next step.
4.20. Repeat steps 4.18-4.19 five more times. A total of six freeze-thaw cycles are required for maximal lysis and extraction. (**Figure 2a,b**)
4.21. Once the freeze-thaw is complete, place samples on ice and *immediately* progress to the next step. Samples should not be frozen at this point for storage as this will diminish available RNA for cDNA preparation.
5. cDNA Synthesis

5.1. Add 2 μL of cDNA synthesis master mix to the PCR tube containing tardigrade lysate. Briefly flick the tube and spin it down at room temperature at 2000 x g for 5 seconds with a tabletop centrifuge before replacing the samples on ice.
5.2. Place the samples in a thermocycler and incubate at 25°C for 10 minutes to anneal primers, at 55°C for 30 minutes to perform reverse transcription, and finally heat inactivate enzymes at 85°C for 5 minutes.
5.3. After the incubation, immediately place the tube on ice and dilute the sample to a total volume of 25 μL by adding 21 μL of sterile nuclease-free water. For low-copy number transcripts, this dilution step can be altered as determined empirically.
6. qPCR

6.1. The annealing temperature of the primer set should be determined using total RNA prepared from larger amounts of tardigrades, for example, the bulk extraction method presented in Boothby, 2018^43^.
6.2. A PCR temperature gradient should be run to determine the optimal annealing temperature before running qRT-PCR (for all PCR settings used in this protocol, refer to **Tables 1 and 2**.)
6.3. Thaw one tube of indicator dye super mix on ice and isolate from light. Place a 96-well qPCR plate on ice and place 5 μL super mix, 2μL of water, 1 μL of each primer (10 μM), and 1μL of cDNA product in the number of desired wells.
6.4. Seal the PCR plate with plate seal and run the qRT-PCR using an annealing temperature appropriate for the primer set. (For all qRT-PCR settings used in this paper, refer to **Table 3**.)
7. Quantification and Results Interpretation

7.1. Results are compared quantitatively to one or more control housekeeping genes, whose expression is expected to be constant over the imposed conditions. For this study, the actin gene was used.
7.2. The C_t_-values or cycle threshold for each well are obtained and compared to the C_t_ values of the control housekeeping gene reactions. The fold change in gene expression is calculated using the following equation:

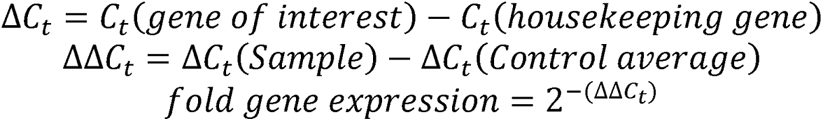

Fold gene expression is plotted for each transcript and tardigrade as a2^-(ΔΔCt)50^.
7.3. To obtain a rough estimate of the transcript number from the C_t_-value, the following equation was used:

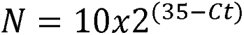

**Figure 1.**
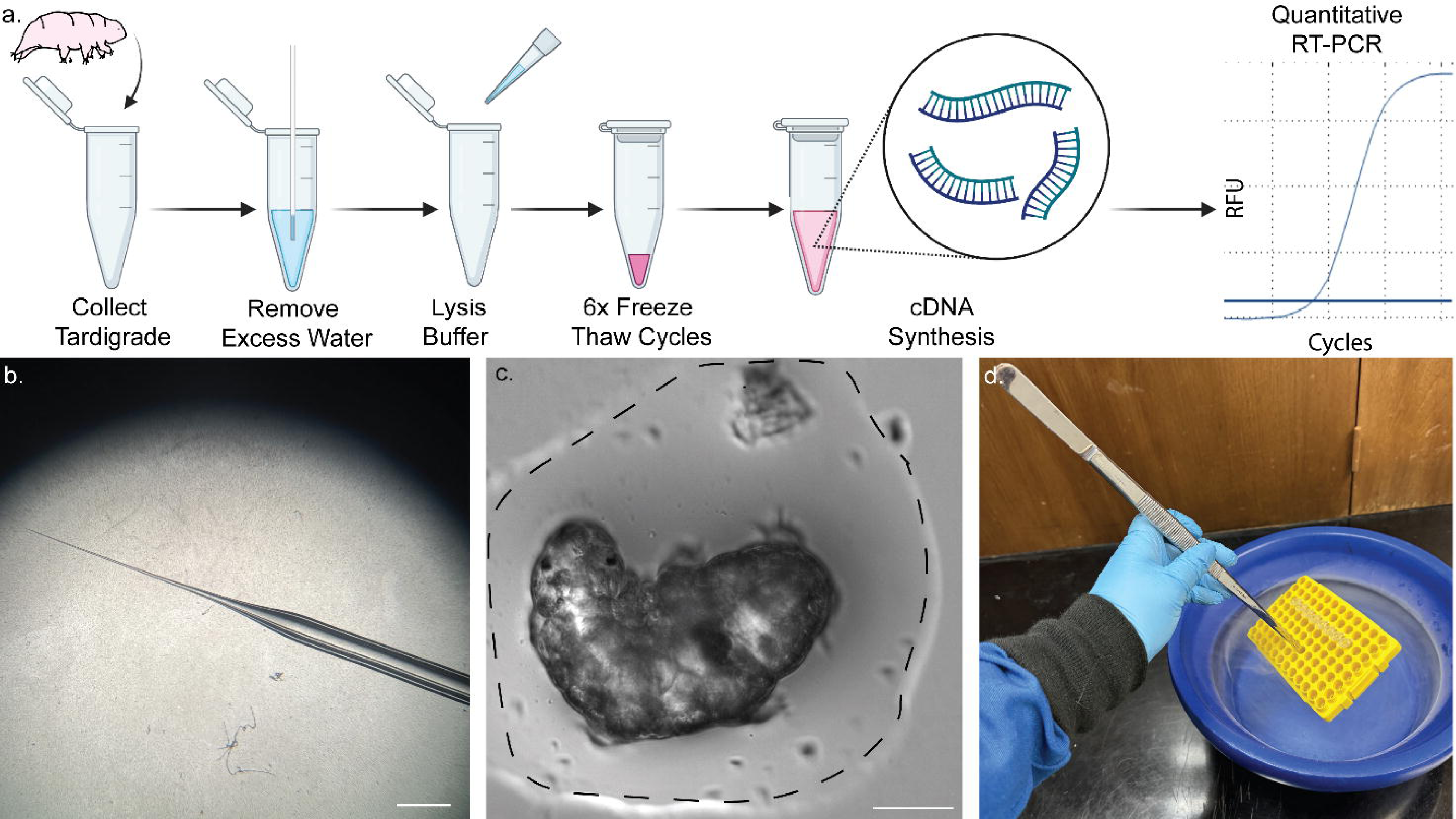
Single-tube pipeline for RNA extraction from a single tardigrade. **a)** Scheme showing the protocol for RNA extraction from a single tardigrade, including six freeze-thaw cycles and subsequent cDNA synthesis. Samples may subsequently be used for RT-PCR and qRT-PCR. **b)** Image of micropipette taper used for removal of water. Scale, 2 mm. **c)** Bright field image of a tardigrade in a small volume of water (dotted line). Removal of most water to the extent shown is required for successful extraction and prevents dilution of lysis buffer. Scale, 50 μm. **d)** Image showing immersion of samples in liquid nitrogen using long forceps to rapidly freeze-thaw the samples safely. Some of the content was created in BioRender. Kirk, M. (2022) BioRender.com/d93s511

**Figure 2:**
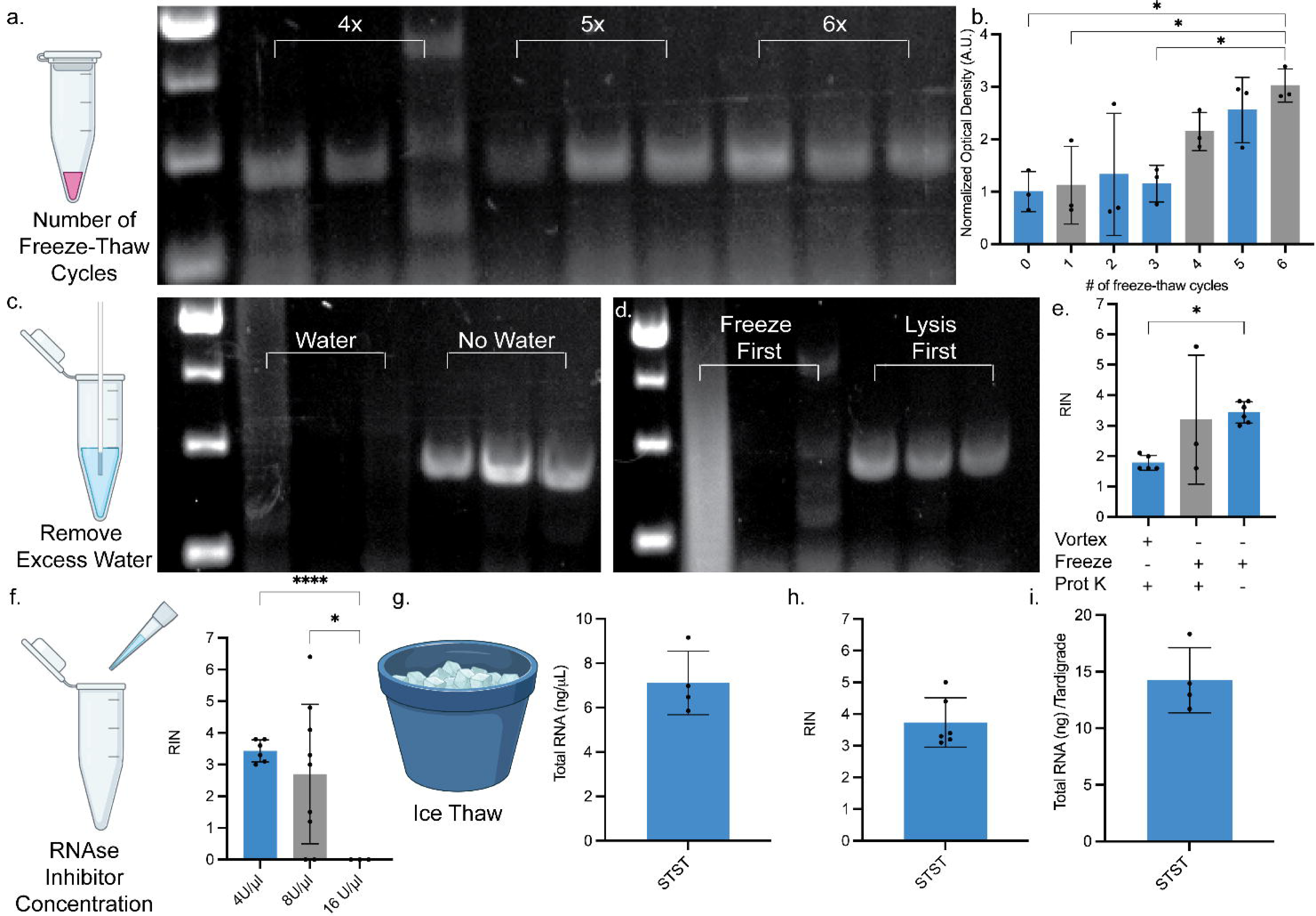
Optimization of single-tardigrade RT-PCR using actin cDNA as a marker for extraction quantity. **a)** Representative gel depicting results from single-tardigrade RT-PCR extracted using 4 (lane 2-4), 5 (lane 5-7), and 6 (lane 8-10) freeze-thaw cycles to enhance lysis after proteinase K treatment and heat-shock. Full gel-containing samples from 1, 2, and 3 freeze-thaw cycles are shown in **Figure S1**. **b)** optical density quantification of ethidium bromide staining of actin RT-PCR across various freeze-thaw cycle numbers. Data represent optical density values of bands from three individual extractions and PCR amplification of actin via RT-PCR per condition. One way ANOVA, with Tukey’s multiple comparisons post hoc 0 vs. 6, p= 0.020 and 3 vs. 6 p=0.022, error bars represent S.D. (Standard deviation) **c)** Representative gel showing the effect residual spring water removal from isolated tardigrades prior to the addition of lysis buffer. Samples containing water (lanes 2-4) and samples where the water was removed (lanes 5-*7*). **d)** Representative gel showing the effect of lysis order, with freeze-thaw performed prior to chemical lysis (lanes 2-4) or chemical lysis with proteinase K prior to freeze-thaw lysis (lanes 5-7). **e)** RNA integrity scores reported by Agilent High Sensitivity Tape station of single tardigrade extracts animals in the absence of freeze-thaw, in the absence of vortexing, and utilizing only freeze-thaws without proteinase k digestion or vortexing. Each data point represents the RIN score from one singular RNA extraction. One way ANOVA, with Tukey’s multiple comparisons post hoc, * p=0.036. **f)** RIN scores reported from single tardigrade extracts in the presence of 4U/μL, 8U/μl and 16U/μL RNAse inhibitor. Brown-Forsythe ANOVA with Dunnett’s T3 multiple comparisons test, **** p=<0.0001 and *=0.0487. **h)** RIN scores from single tardigrade extracts thawing on ice. **i)** RNA quantity in ng/μL as measured using Qubit high sensitivity RNA kit. **j)** RNA quantity in ng per single tardigrade extract using STST. 5 μL of PCR products were loaded per lane unless otherwise noted. All error bars are reported S.D.. Some of the content was created in BioRender. Kirk, M. (2022) BioRender.com/d93s51

**Table 1.** PCR thermocycling protocol for non-quantitative PCR. The table describes the exact thermocycling procedure used for this study. Tm is denoted as the melting temperature for a reaction. The specific melting temperatures for each reaction can be found in **Table 2**.

| Step | Temp. (°C) | Duration | No. of cycles |
| --- | --- | --- | --- |
| Initial denaturation | 95 | 60 s | 1 |
| Denaturation | 95 | 15 s | 35 |
| Annealing | Tm | 30 s |  |
| Extension | 72 | 1 min |  |
| Final Extension | 72 | 5 min | 1 |
| Hold | 4 | ∞ | - |

**Table 2.**
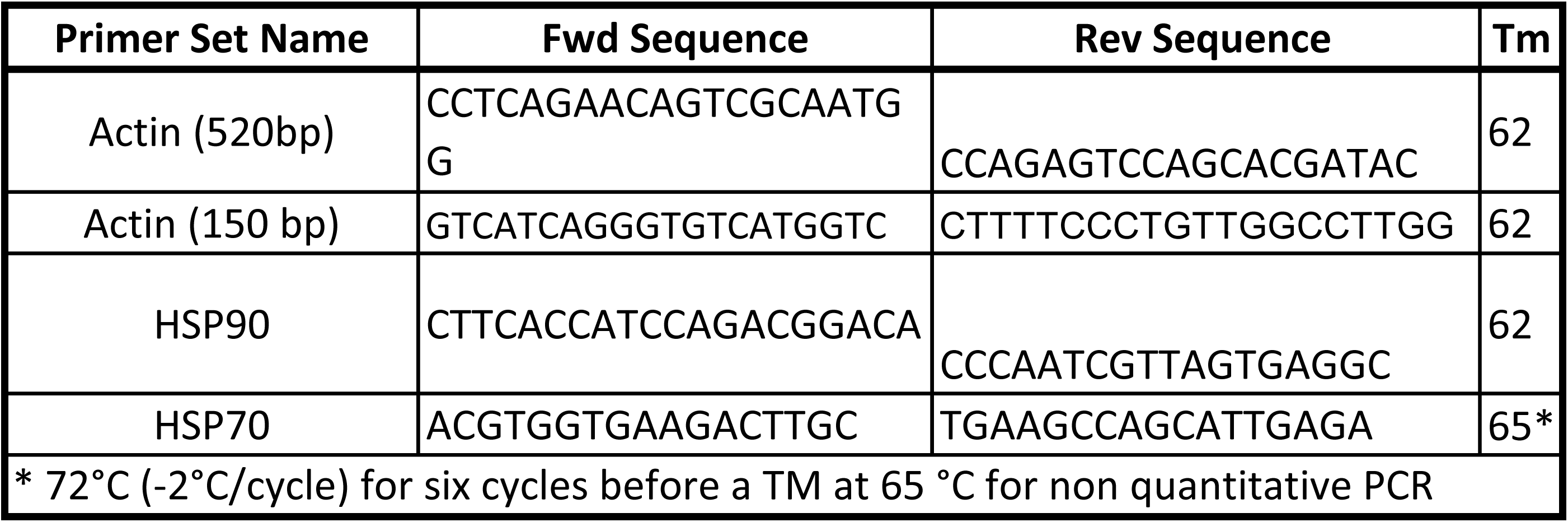
Melting temperature and sequences for primers used in this study. The table shows the forward and reverse sequences of primers used for this study and their optimized melting temperature. It is important to note that HSP70 required a brief touch-down protocol consisting of 6 cycles before the amplification to ensure specificity.

**Table 3.** Thermocycling conditions for qPCR. The table depicts the thermocycling settings for qPCR reactions used in this study. The melting temperatures or Tm’s were identical to those shown in **Table 2**.

| Step | Temp. (°C) | Duration | No. of cycles |
| --- | --- | --- | --- |
| Initial denaturation | 95 | 60 s | 1 |
| Denaturation | 95 | 15 s | 30 |
| Annealing Extension | Tm | 30 s |  |
| Denaturation | 95 | 10 s | 1 |
| Melt Curve Analysis | 65 (+0.5°C) | 10 s | 60 |

Where N is the number of transcripts, and 2 is the assumed PCR efficiency or the fold increase in fluorescence per cycle of PCR^50^.

## RESULTS

### Development and optimization of single-tardigrade RNA extraction

Adapting the protocol from Ly et al., 2015^45^ for RNA extraction in tardigrades, the STST system is optimized to maximize the quantity and quality of the preparation (**Figure 1a**). RT-PCR was performed for actin transcripts, quantifying transcript yield by amplifying a 527 bp region spanning exons 1 and 2. (Sequences for these primers can be found in **Table 1**). The optical density of the expected actin band was quantified as fluorescence intensity with ImageJ/FIJI. All regions of interest quantified were of equal areal size across each gel.

First, methods of mechanical lysis required for disrupting the cuticle of the tardigrade were assessed. In contrast to a single-animal RNA extraction protocol described for *C. elegans*^45^ which reported that Proteinase K lysis and a heat-shock were sufficient to lyse the animals, tardigrade extraction required a minimum of six freeze-thaw cycles to achieve consistent and robust RNA extraction (**Figure 2a, Figure S1**).

These samples were vortexed at 2700 rotations per minute for 15 seconds between each freeze-thaw cycle to promote cell lysis. To minimize time and potential degradation, the freeze-thaw cycle number was kept at this minimal value throughout the remainder of the preparations. At lower numbers of freeze-thaw cycles (1-5), higher levels of genomic DNA (gDNA) products, as evidenced by the appearance of slower-migrating bands than expected for the actin cDNA, were found. This suggests that the freeze-thaw process prior to dsDNA removal by DNase is required for the complete removal of gDNA. The lower number of freeze-thaw cycles may decrease the probability that all nuclei are effectively permeabilized, thus preventing dsDNase from degrading all gDNA during the 25°C incubation period. At higher temperatures, during the reverse transcription and heat inactivation steps, the remaining nuclear DNA may be released and serves as a template during subsequent PCR. (**Figure 2a, Figure S1**). Ultimately, the consistency of extraction success and the yield, as measured by optical density, increased with higher numbers of freeze-thaw cycles, with six cycles showing a three-fold increase in cDNA yield compared to no freeze-thaw cycling (**Figure 2b**).

Next, we tested whether the near-complete removal of excess water carried over from the sample transfer was essential to achieve consistent lysis of single tardigrades. The STST protocol uses a minimal volume of lysis buffer to maximize RNA concentration. We were concerned that excess water might dilute the detergent and EDTA in the lysis buffer and thus interfere with reliable lysis. **Figure 2c and Figure S2** show the results of triplicate individual tardigrade extractions in the presence and absence of residual water from the tardigrade transfer. These data indicate that removing excess water is critical to success, as product was not observed in samples containing excess water.

We sought to determine whether the order of lysis is important for achieving robust extraction of RNA and found that performing the freeze-thaw step before the Proteinase K and enzymatic lysis resulted in little or no detectable RNA by RT-PCR (lanes 2-4 of **Figure 2d**). However, PCR products were readily obtained when chemical lysis was performed before the freeze-thaw step (lanes 5-7 of **Figure 2d**). This suggests either that Proteinase K digestion of the cuticle is required before mechanical lysis or that prior freeze-thaw lysis extracts the RNA, leaving it exposed to endogenous RNAse activity throughout the proteinase K treatment.

To assess whether proteinase K digestion is required for the isolation of RNA from single tardigrades, RNA quality was assessed via high-sensitivity tape station. Animals that experienced vortexing after proteinase K treatment showed high RNA fractionation, while freeze-thaw cycling after proteinase K treatment resulted in increased yet inconsistent RNA integrity scores (RIN) (**Figure 2e).** Intriguingly, RNA extraction in the absence of the proteinase K and vortexing steps increased consistency and quality significantly, suggesting that the proteinase K treatment is neither required nor helpful for RNA extraction from *H. exemplaris* (**Figure 2e).** Eliminating this treatment significantly reduced the time required for our protocol from 45 minutes with proteinase K treatment to 7 minutes in our improved STST protocol. Further, increasing the RNAse inhibitor concentration from 4U/μL to 8U/μl or 16 U/μL resulted in incomplete freezing of the animals, yielding inconsistent results at 8U/μL and a complete lack of freezing and subsequent lysis at 16 U/μL (**Figure 2f**). This effect is most likely attributable to the RNAse storage solution containing 50% glycerol, which acts as a cryoprotectant. Also, we found that thawing on ice resulted in moderately increased RIN scores (**Figure 2g).**

Finally, to assess the total yield of STST preparations, RNA quantity per μL was measured via High Sensitivity RNA Qubit. This analysis revealed a yield of 7.11 +/- 1.43 ng/μL (**Figure 2h**), suggesting that the STST preparation yields 14.24 +/- 2.88 ng per single *Hypsibius exemplaris* animal (containing ∼1400 cells) (**Figure 2i**). This result is consistent with the yields observed with *C. elegans* adult (∼3000 cells, including germ cells), which yield ∼ 35 ng per animal^45^.

### Comparison to existing tardigrade RNA extraction protocols

In developing the STST RNA extraction protocol, we aimed to limit the time required to perform the technique, the number of sample-experimenter interactions, the cost, and the number of animals required for robust RNA extraction. **Figures 3a** and **3b** depict a schematic contrasting a previously described single-tardigrade RNA extraction protocol that has been used for RNA sequencing following linear PCR amplification (**Figure 3a**) and the single-tube, single-tardigrade RNA extraction method (**Figure 3b**), highlighting each method in terms of these four parameters. Previously, single-tardigrade RNA extraction protocols required approximately ten minutes, three tubes, and five experimenter interactions before cDNA synthesis; however, these methods have not yet been assessed for qRT-PCR applications on single animals. Current published RNA extraction protocols for qRT-PCR assessment of transcriptional changes use a minimum of seven tardigrades^24^. The STST method described here decreases the time to seven minutes with only one sample-experimenter interaction and a single tube. Removing sample-experimenter interactions reduces the potential for RNAse contamination, thus increasing the potential for high-quality extractions. ^24^

**Figure 3.**
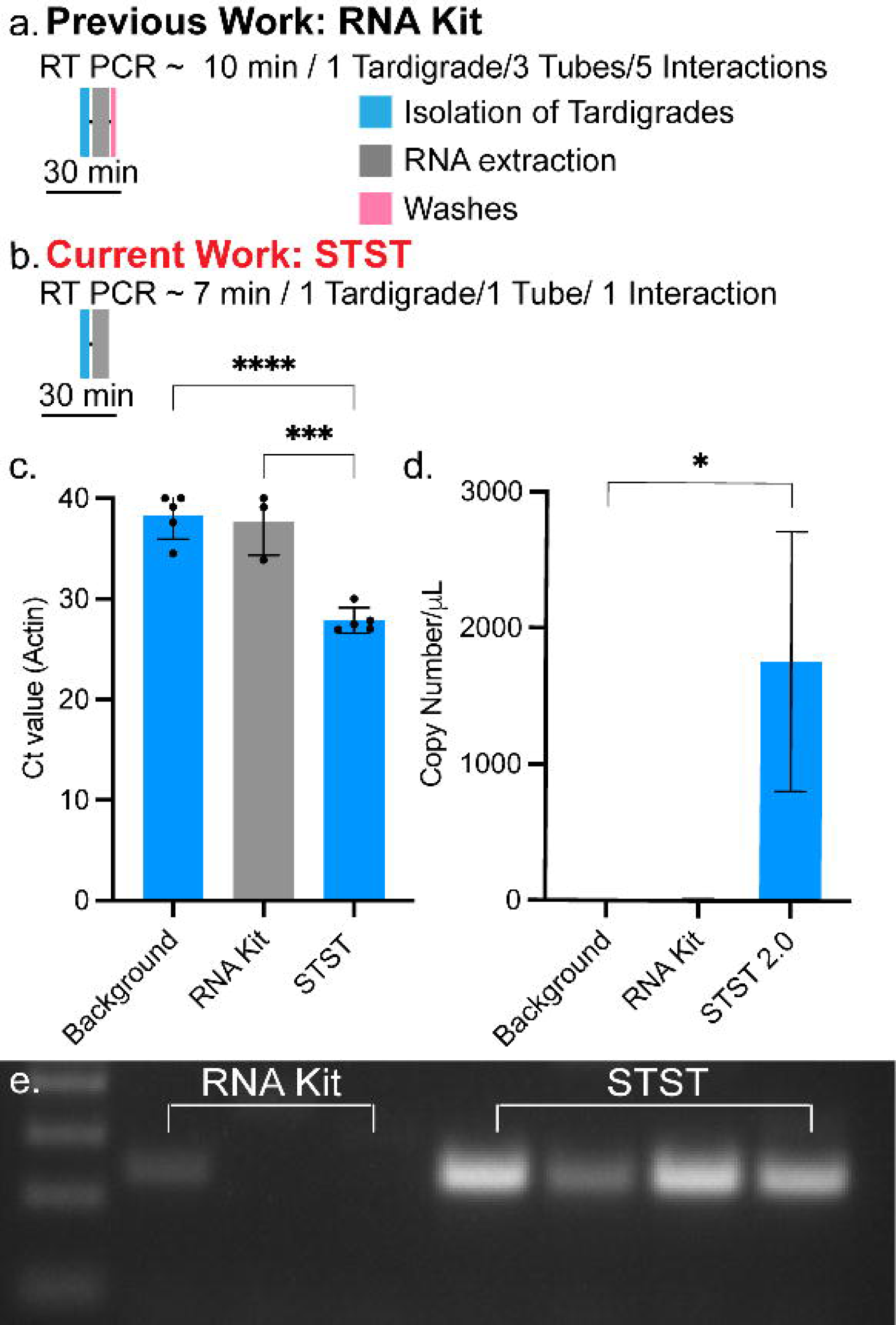
Comparison of RNA extraction protocols. Schematized time-course for **a)** an existing phenol and guanidine isothiocyanate-based RNA extraction kit, requiring ∼10 minutes for extraction (Scale bar is 30 minutes) and **b)** The single-tube, single-tardigrade protocol that permits extraction in 7 minutes and requires one tube. The time courses are drawn to scale. Scale bar is 30 minutes. **c)** Graph showing C_t_ values of actin qRT-PCR reactions run in triplicate where each data point represents the average Ct values of three technical replicates from one individual tardigrade extract. Single-tardigrade-based extractions and the background control revealing background fluorescence from samples run on qRT-PCR in the absence of template. One way ANOVA, Tukey’s multiple comparisons test ***, p=0.0003 and ****, p<0.0001, error bars reflect S.D. **d)** Estimated transcripts per μL of sample derived from the C_t_ values depicted in **(c)**.Kruskal-Wallis Test, Dunn’s multiple comparisons test, *, p=0.0165 error bars reflect S.D. **e)** Representative gel showing actin RT-PCR from single-tardigrade cDNA samples extracted using RNA kit (lanes 2-4) and STST lanes (5-8). Gels were loaded with 6 μL of sample and 1 kb plus DNA Ladder

To compare the efficacy of STST extractions with the RNA extraction kit, single-tardigrade extractions were performed using both methods, and transcript numbers were quantitatively assessed using qPCR. Amplifying a 150 bp region of the housekeeping gene encoding actin, we used the C_t_ value, i.e., the cycle number required for the fluorescence signal to be detected above the background, to indicate cDNA quantity. The average C_t_ value of single tardigrades processed with the STST protocol was 27.86 +/- 1.268 (**Figure 3c**), while those extracted with the RNA extraction kit were 37.67+/- 3.311 (**Figure 3c).** While both protocols yield a signal above background fluorescence obtained from control qPCRs run without template cDNA, only the STST protocol was statistically significant compared to background levels. This finding indicates that although RNA was present in the RNA extraction kit samples, the kit did not produce sufficient quantities of RNA to be consistently detected. Using these C_t_ values, we were able to estimate the transcript number per μL of cDNA: STST yielded ∼1760+/-954.2 actin transcripts per μL, while the RNA Kit-based method yielded ∼ 7.5+/- 12.29 actin transcripts per μL (**Figure 3d),** suggesting that as a single-tardigrade extraction method, the STST system is > 200-fold more efficient. When quantified, the STST protocol produced significantly more transcripts than the RNA extraction kit, which was again statistically indistinguishable from background measurements (**Figure 3d).** This suggests that the current RNA kit-based method for single tardigrades does not sufficiently extract RNA or loses significant amounts of RNA throughout the purification process, resulting in undetectable quantities of cDNA downstream. To visualize this effect, qPCR samples were run on an analytical gel, confirming minimal amplification of RNA kit-extracted products (**Figure 3e**, lanes 2-4) and robust amplification in the STST-extracted samples (**Figure 3e**, lanes 5-8).

Finally, we compared the cost of RNA extraction and cDNA synthesis using the RNA extraction kit with the STST method (**Table 4**). As the elution volume for the RNA kit was three times that of the STST protocol, it requires three times the cDNA synthesis material, accounting for the higher cost in cDNA synthesis for each whole tardigrade extract before samples were diluted to equal volumes of 25μL. Further, as we had previously determined (**Figure 3 c and d**) that a linear amplification step would be required to yield quantities above background C_t_ values, we have included that step in our cost estimates as directly described by Arakawa et al. 2016^44^; however, there may be a more economical way to perform linear amplification of which we are unaware.

**Table 4:**
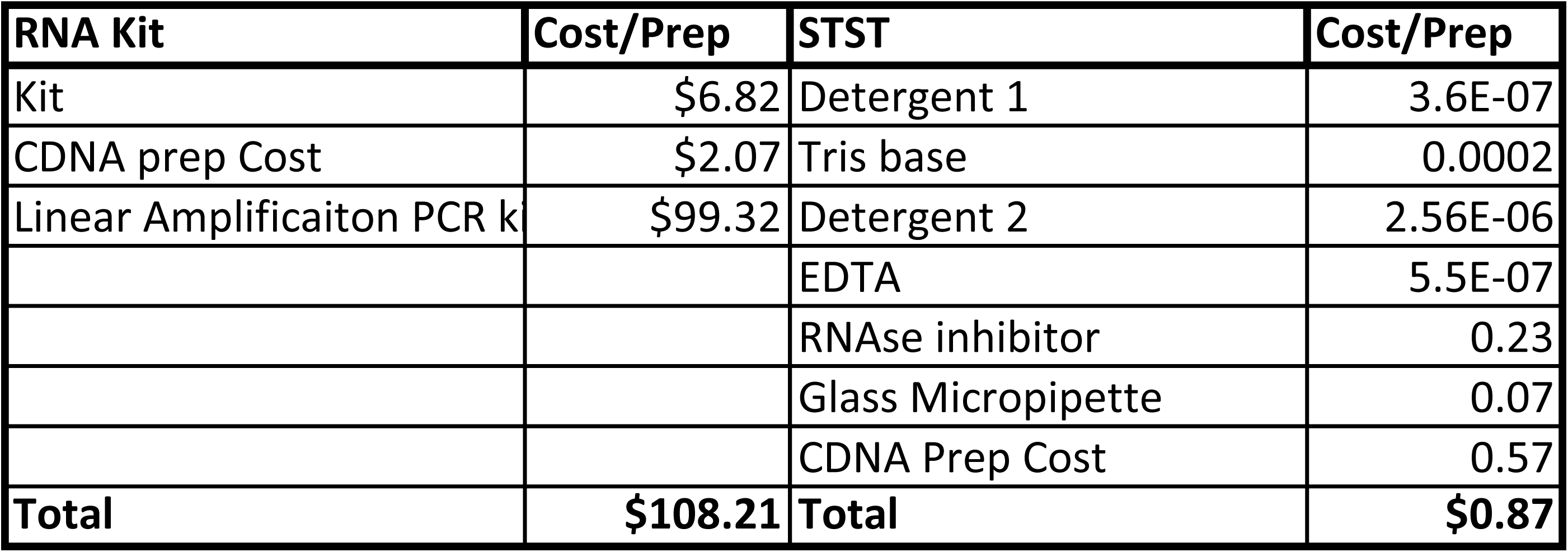
Cost breakdown for RNA extraction. The table describes the cost breakdown for each RNA extraction method per single animal. Prices were obtained from the vendor website on September 30^th^, 2024, and are subject to change.

### Application of the single-tube single-tardigrade extraction protocol for quantifying heat-shock responses in H. exemplaris

We next sought to explore whether this protocol can be applied to effectively monitor transcriptional changes in *H. exemplaris*, turning to the canonical heat-shock response pathway, which results in the upregulation of both HSP70β2 and HSP90α (**Figure 4a**) following brief exposure to high temperatures^45^. Tardigrades in their active state are highly sensitive to increases in temperature^29^; thus, we selected 35°C for 20 minutes as the heat-shock, as we found that this resulted in minimal lethality (**Figure 4b**). We observed that the response follows a similar pattern to that seen in other organisms^45^. We first exposed tardigrades to either rearing temperature (23°C) or high heat (35°C) for 20 minutes (**Figure 4c**). We then extracted RNA from individual animals, and generated cDNA using the STST method by harvesting immediately or at 1 hour, 2 hours, 4 hours, and 6 hours after exposure to the heat-shock. We subsequently quantified transcripts for HSP70β2 (**Figure 4d**) and HSP90α (**Figure 4e**) using qRT-PCR. Significant upregulation of HSP70β2 (∼11-fold) and HSP90α(∼4-fold) was observed after 1 hour, and expression slowly returned to baseline over the next several hours. These results follow the expected patterns of heat-shock responses seen in *C. elegans*^45^.

**Figure 4.**
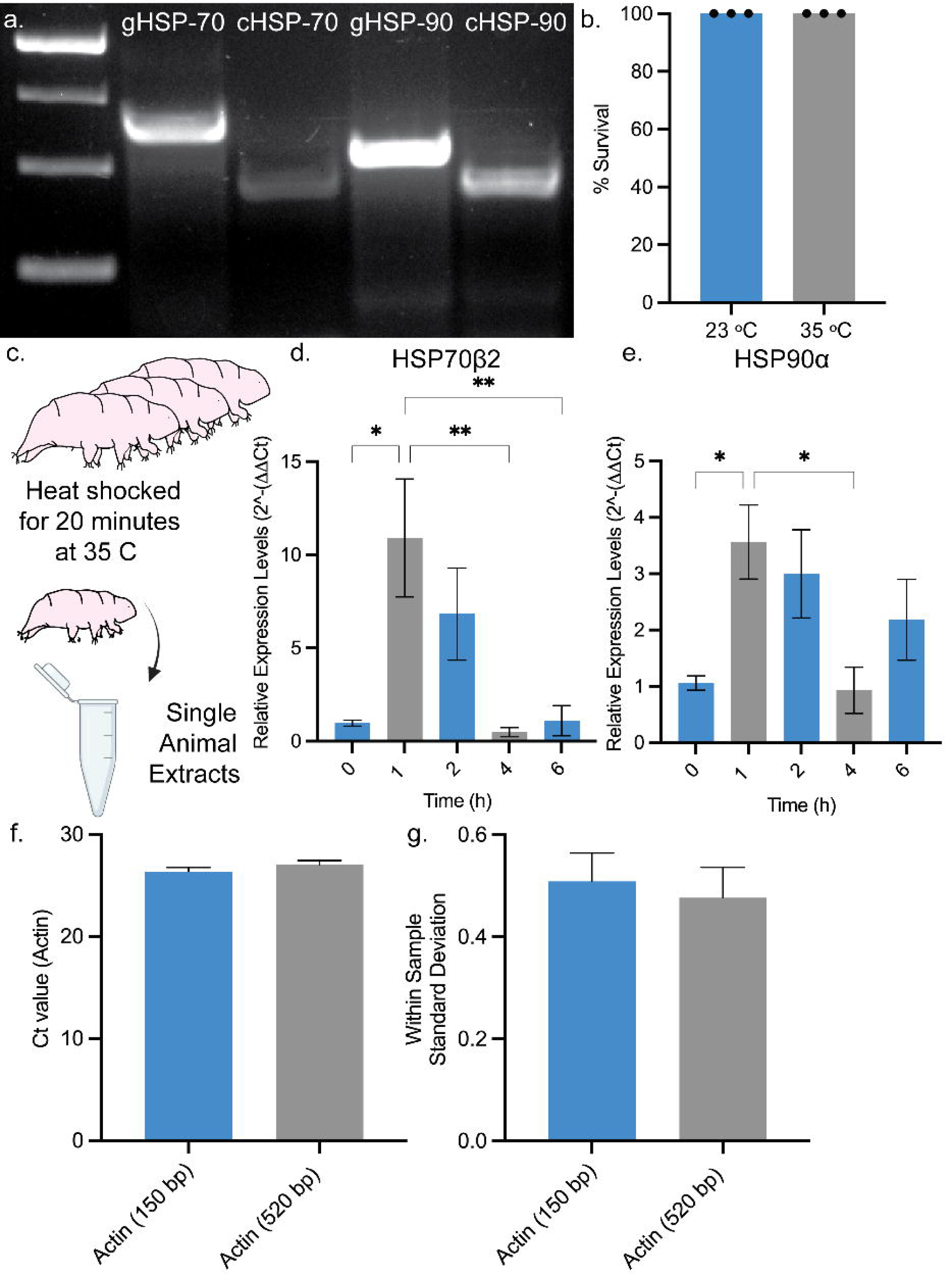
Expression of HSP-70 and HSP-90 transcripts. **a)** Representative gel showing amplification of HSP70 from genomic DNA (gDNA; lane 2) and cDNA (lane 3) and HSP90 from gDNA (lane 4) and cDNA (lane 5). Variance in gDNA and cDNA lengths reflects the primer design, which spans an intron. **b)** Survival of tardigrade post exposure to 23°C and 35°C for 20 minutes. Each data point represents percent survival from a cohort of six animals. T-test found no statistical difference**. c)** Conditions analyzed. **d)** Relative expression levels of HSP70β2 at the indicated time post-20-minute heat-shock at 35 °C. Kruskals-Wallis Test with Dunn’s multiple comparisons analysis *, p=0.0339 (1 vs. 0)**, p=0.0043 (1 vs. 4), **, p=0.0097 (1 vs. 6). Each data point represents normalized expression levels from an individual tardigrade RNA extract averaged across three technical replicates. **e)** Relative expression levels of HSP90α at the indicated time post-20-minute 35°C heat-shock. Brown-Forsythe ANOVA with Dunnett’s T3 multiple comparisons test*, p=0.0382 (1 vs. 0) *, p=0.0421 (1 vs. 4). Each data point represents normalized expression levels from an individual tardigrade RNA extract averaged across three technical replicates. **f)** Raw C_t_ values for actin qRT-PCR amplifying either a 150 bp or 527 bp amplicon from all samples presented in panels **d** and **e**. Each data point represents C_t_ from an individual tardigrade RNA extract averaged across three technical replicates; T-test found no statistical difference. **g)** Standard deviation in C_t_ values across technical replicates for each extract represented in panels **d** and **e.** Reactions amplified an actin amplicon of either 150 bp or 527 bp; T-test found no statistical difference. Some content was created in BioRender. Kirk, M. (2022) BioRender.com/d93s51

Finally, to assess the quality of STST as an extraction method for qRT-PCR, we compiled all actin C_t_ values across all the samples run for panels **4d** and **e.** The average actin C_t_ values for an actin 150 bp amplicon and an actin 527 bp amplicon were not statistically different (**Figure 4f**). Furthermore, the comparison of variance (standard deviation) across C_t_ values derived from triplicates obtained from each individual extract were consistently below the average quality cut-off of 0.5 cycles per individual extract^50^ (**Figure 4g**).

## DISCUSSION

This study presents an efficient method for the extraction of RNA for single-tardigrade qRT-PCR. Directly comparing the STST methodology to an existing single tardigrade RNA extraction kit revealed that STST RNA extraction yields >200-fold higher amounts of actin RNA transcripts, reduces the cost to less than one dollar per sample, and reduces the time required for extraction by 30%. To apply STST to a relevant biological question, we assessed the short-term heat-shock response expression profile. We found that transcripts for both HSP70β2 and HSP90α, as expected, were strongly upregulated 1 hour after heat exposure.

Although the STST protocol is a substantial improvement over previous methods, several limitations present opportunities for improvement and further assessment. First, we have not evaluated the ability of STST to detect transcripts expressed at very low levels. Troubleshooting for appropriate template concentration may be required for transcripts which are expressed at low levels. Secondly, while the STST method showed an ∼ 80% success rate, defined as the percent of extracts resulting in an actin C_t_ values less than 31, there were occasions when the method failed to obtain usable quantities of cDNA, perhaps because of RNAse activity or insufficient lysis. Optimizing the inhibition of RNAse activity might help increase the success rate. Finally, RNA quality could be improved in this method. As in this study, measures can be taken to circumvent the effects of degradation and subsequent fragmentation of the RNA, as well as to mitigate its impact on qRT-PCR results. Improvements in RNA quality would further enhance the rapid assessment of RNA transcriptional changes, the analysis of RNAi knockdown efficacy, and the ability to evaluate variance across populations of many individual animals. It is important to note that all protocols for RNA extraction in tardigrades require some form of mechanical lysis. Traditional phenol and guanidine isothiocyanate protocols require three freeze-thaw cycles^43^, while the RNA extraction kit requires manual rupturing with a pipette^36, 44^ Ultimately, different methods of mechanical lysis may improve quality in the future.

We identified several critically important steps in the STST method. First, removing most of the water from the tube prior to adding lysis buffer is critical: we leave a bubble approximately two tardigrade lengths in diameter surrounding the animal. Though we have not measured this minute volume, removing as much water as possible will improve consistency and overall success (Figure 1c). Second, the immediate transfer of the samples into a cDNA preparation reaction is important. Our preparation does not include any RNA purification, so it is imperative that one does not pause the procedure after extraction. It is possible that extracts could be successfully stored at −80 °C; however, we have not assessed the effect of such storage on qRT-PCR quality. Third, adhering to the guidelines for performing qRT-PCR on mildly degraded samples is vital, i.e. analyzing short amplicons of < 200 bp using multiple housekeeping genes to normalize data whenever possible^51, 52^.

Overall, our findings represent an important step in the growing repertoire of molecular tools available in tardigrade research. This method complements existing RNA extraction methods^43, 44^, which have provided techniques for single tardigrade quantification of transcriptional changes via RNA-seq following a linear amplification step^43, 44^. STST allows for quantification of transcripts isolated from single tardigrades without the need for a scale-up step. As most tardigrade responses to extreme conditions, such as those triggered by desiccation, result in ∼75-80% survival rates in *H. exemplaris*; bulk samples contain mixed populations of tardigrades, including both survivors and non-survivors. Such a mixed population has the potential to confound the analysis of transcriptional changes that occur during response and recovery from extreme stresses. In bypassing the need for bulk preparation, STST and the tardigrade RNA-seq method^36, 44^ avoid the confounding problems inherent in a mixed population and allow for the full dynamic range of a response to be quantified. We envision that STST can be used not only to quantify the range of transcriptional responses and assess the efficacy of RNAi knockdowns, but also be used in many other applications in the future.

## DATA AVAILABILITY

All raw analytical gel data has been incorporated into the supplemental data of this manuscript. Transcript sequences were identified via Ensembl search at: https://metazoa.ensembl.org/Hypsibius_exemplaris_gca002082055v1/Info/Index?db=core HSP70β2 (BV898_04401), HSP90α (BV898_50798) and Actin(BV898_02877).

## ACKNOWLEDGMENTS

We would like to acknowledge the NIH Ruth Kirschstein Fellowship # 5F32AG081056-02 and the Errett Fisher Post-Doctoral Fellowship, which supported Dr. Molly J. Kirk, the Crowe Family Fellowship, which supported Chaoming Xu, and a University of California, Santa Barbara Academic Senate Grant, and NIH grants # R01GM143771 and #2R01HD081266, which supported these research efforts. The authors also acknowledge the use of the Biological Nanostructures Laboratory within the California NanoSystems Institute, supported by the University of California, Santa Barbara and the University of California, Office of the President.

---

| Name of Material/ Equipment | Company | Catalog Number | Comments/Description |
| --- | --- | --- | --- |
| Corning™ PYREX™ Reusable Media Storage Bottles | Thermo Fisher Scientific | 06-414-1E | Referred to as 2 L Autoclave-safe Glass Bottle |
| Andwin Scientific 3M LEAD FREE AUTOCLAVE TAPE 1" | Thermo Fisher Scientific | NC0802040 | Referred to as Autoclave Tape |
| *Hypsibius exemplaris* Z151 Strain | Carolina | 133960 | Referred to as Tardigrades or *H. exemplaris* |
| *Chlorococcum hypnosporum* . | Carolina | 152091 | Referred to as Algae |
| Direct-zol Micro Prep | Zymo Research | R2060 | Referred to as RNA extraction kit |
| RNasin® Ribonuclease Inhibitor | Promega | N2111 | Referred to as RNase inhibitor |
| Maxima H Minus First Strand cDNA Synthesis Kit | Thermo Fisher Scientific | K1651 | Referred to as cDNA Synthesis Master Mix |
| SsoAdvanced™ Universal SYBR® Green Supermix | BIO RAD | 1725271 | Referred to as Indicator Dye Super mix |
| Triton™ X-100 | Fluka | 93443 | Referred to as Detergent 1 |
| Tris base | Fisher Scientific | T395-500 | Referred to as Tris or Tris Base |
| TWEEN® 20 | Sigma aldrich | P1379-500 | Referred to as Detergent 2 |
| EDTA | Fisher Scientific | S311-500 | Referred to as EDTA |
| Water - PCR/RT-PCR certified, nuclease-free | Growcells | PCPW-0500 | Referred to as Sterile Nuclease Free Water |
| Spring water | Nestle Pure Life | 44221229 | Referred to as Spring Water |
| RNase-Free, 8 Strip 0.2mL PCR Tubes with caps | Invitrogen | AM12230 | Referred to as Sterile PCR Tube |
| BioRadHard-Shell® 96-Well PCR Plate | BioRad | HSS9641 | Referred to as PCR Plate |
| PETRI DISH, PS, 35/10 MM, WITH VENTS | Grenier | 627102 | Referred to as 35 mm Petri dish |
| Thermo Scientific™ Nunc™ Petri Dishes | Thermo Fisher Scientific | 08-757-099 | Referred to as 100 mm Petri dish |
| Supertek Scientific Tirrill Burners | Thermo Fisher Scientific | S09572B | Referred to as Bunsen Burner |
| Dumont 5 Biology Tweezers | Fine Science Tools | 11254-20 | Referred to as Fine Forceps |
| Fisherbrand™ Economy Impact Goggles | Fisher Scientific | 19-181-501 | Referred to as Splash Goggles |
| Nitrile Gloves | Fisher Scientific | 17-000-314 | Referred to as Nitrile Gloves |
| BULWARK FR Lab Coat: | Grainger | 26CF64 | Referred to as Lab Coat |
| Tempshield™ Cryo-Gloves™ | Fisher Scientific | 11-394-305 | Referred to as Cryo Gloves |
| PIPETMAN P1000, 100-1000 µL, Metal Ejector | Gilson | F144059M | Referred to as P 1000 Pipette |
| PIPETMAN P200, 20-200 µL, Metal Ejector | Gilson | F144058M | Referred to as P 200 Pipette |
| PIPETMAN P10, 1-10 µL, Metal Ejector | Gilson | F144055M | Referred to as P 10 Pipette |
| Stereo-Microscope System w/optics and illumination | TriTech Research | SMT1 | Referred to as Dissecting Microscope |
| Autoclave Tape | Thermo Fisher Scientific | AB1170 | Referred to as PCR Plate Seals |
| Pound This 4-Color Modeling Clay | American Science Surplus | 96517P001 | Referred to as Clay |
| C1000 Touch™ Bio-rad Thermocycler | BioRad | 1851148 | Referred to as Thermocycler |
| C1000 Touch™ Bio-rad Thermocycler with CFX™ Optics Module | BioRad | 1845097 | Referred to as qPCR thermocycler |
| Glass Micropipette O.D. 1mm ID 0.58, Length 10 cm | TriTech Research | GD-1 | Referred to as glass micropipette |
| Narishige Dual-Stage Glass Micropipette Puller | Tritech Research | PC-10 | Referred to as micropipette puller |
| Filament for pipette Puller | Tritech Research | PC-10H | Referred to as Filament |
| Table Top Centrifuge | Qualitron | DW-41-115-NEW | Referred to as Table Top Centrifuge |
| Liquid Nitrogen Dewar 1L | Agar Scientific | AGB7475 | Referred to as Cryo-safe container |
| Daigger & Company Vortex-Genie 2 Laboratory Mixer | Thermo Fisher Scientific | 3030A | Referred to as Vortexer |
| 4 Star Straight Strong Medium Point Tweezer | Excelta | 00-SA-DC | Referred to as Long forceps |
| 96-Well PCR Rack with Lid Assorted, 5 Racks/Unit | Genesee Scientific | 27-202A | Referred to as PCR Rack |
| Prism v10.0 | GraphPad | N/A | Referred to a Prism |
| Benchling v8 | Benchling | N/A | Referred to as Benchling |
| FIJI v 2.14.0/1.54f | ImageJ, | N/A | Referred to as FIJI/ImageJ |
| 1000µl Premium Pipet Tips, Low Binding, Racked, Sterile | Genesee Scientific | 23-165RS | Referred to as Sterile Filter-Tipped P 1000 Pipette Tips |
| 200µl Premium Barrier Tips Low Binding, Racked, Sterile | Genesee Scientific | 23-412 | Referred to as Sterile Filter-Tipped P 200 Pipette Tips |
| 10µl Premium Barrier Tips Low Binding, Racked, Sterile | Genesee Scientific | 23-401 | Referred to as Sterile Filter-Tipped P 10 Pipette Tips |

