## Supplemental Figures for "Single-animal, single-tube RNA extraction for comparison of relative transcript levels via qRT-PCR in the tardigrade *Hypsibius exemplaris*"

DOI:

**Figure S1**

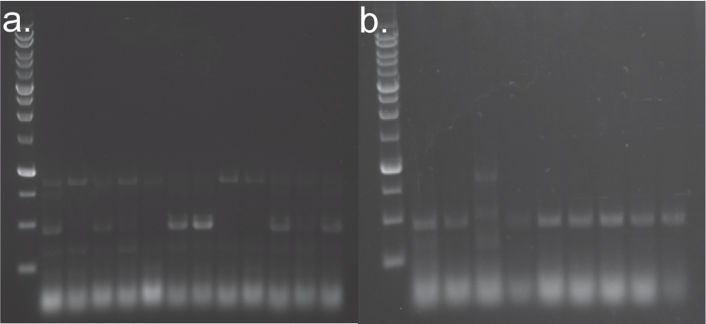

**Figure S1. Freeze-thaw cycle number assessment.**

Agarose gel electrophoresis of actin PCR products from cDNA samples that underwent different numbers of freeze-vortex cycles. **a)** Actin transcript amplification through PCR performed with samples that have been freeze-vortexed for 0-3 times (n=3; 0 cycle: 2-4 lane; 1 cycle:5-7 lane; 2 cycles: 8-10 lanes; 3 cycles: 11-13 lane). 1 kb plus DNA ladder is shown in the first lane; **b)** Actin transcript amplification by PCR performed with samples that have been freeze-vortexed 4-6 times (n=3; 4 cycle: 0-3 lane; 5 cycles: 4-6 lane; 6 cycles: 7-9 lanes) 5 μL of 1 kb plus DNA ladder was loaded in the first lane.

**Figure S2**

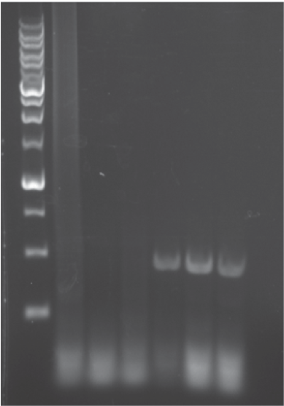

**Figure S2. Water removal is required for successful RNA extraction**

A gel image shows the actin amplicons from PCR performed on samples without (lane 2-4) or with (lane 5-7) water removed using a pulled glass micropipette before adding the lysis buffer (n=3). The first lane was loaded with 5 μL of 1 kb plus DNA ladder, and all gels were loaded with 5 μL of PCR products.

**Figure S3**

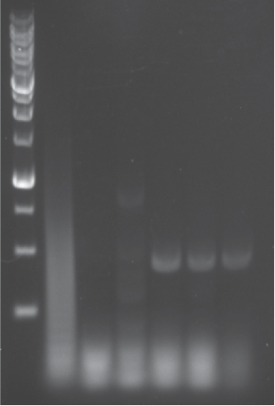

**Figure S3. Chemical and physical lysis order determines extraction quality.**

Gel electrophoresis of actin PCR products from samples that underwent freeze-vortex first (Lane 2-4) versus chemical lysis first (Lane 5-7). The first lane was loaded with 5 μL of 1 kb plus DNA ladder.

**Figure S4**

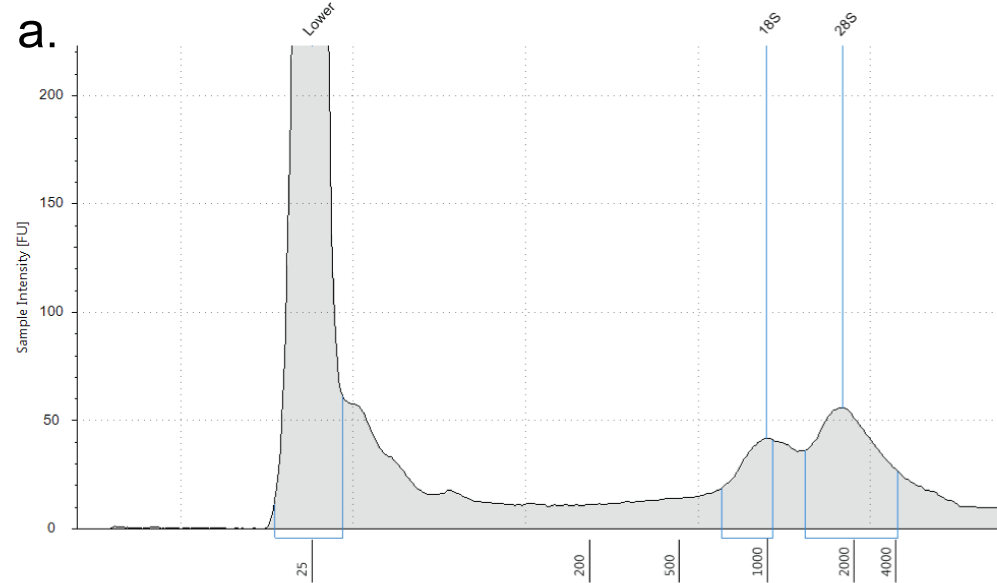

**Figure S4: RNA tape stations run on single tardigrade extracts.** Panel **a)** example tape station run on RNA extracted from single tardigrades using the STST method.

**Figure S5**

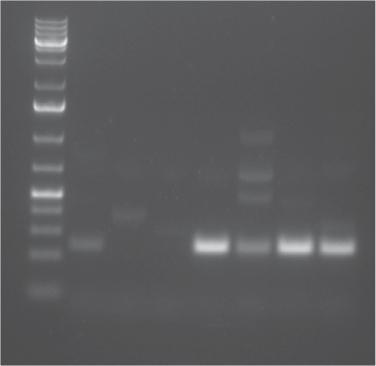

**Figure S5. Comparative analysis of single-tardigrade extraction using STST vs. RNA extraction kit.** Actin PCR products obtained from single-tardigrade cDNA samples extracted using the RNA extraction protocol from Arakawa, 2018 (lane 2-4) versus extraction using the STST method (lane 5-8). The first lane was loaded with 6 μL of 1 kb plus DNA ladder. All electrophoresis was performed with 6 μL of PCR products loaded in a single lane.

**Figure S6**

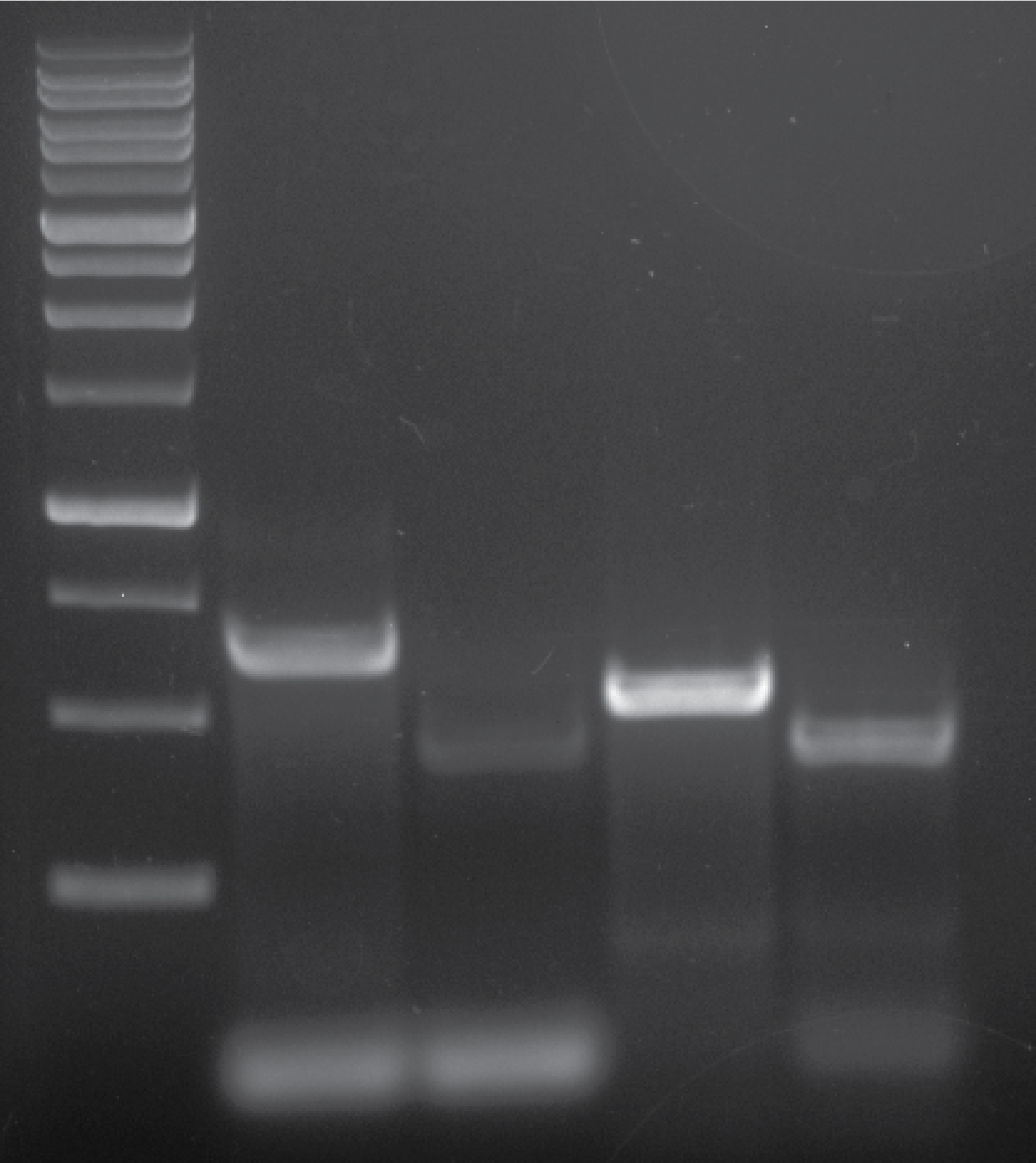

**Figure S6**. Gel electrophoresis of HSP70 and HSP90 amplicons. PCR products from cDNA and gDNA using the He_HSP70B2 and He_HSP90ALPHA primer sets. HSP70B2 amplicon from gDNA is shown in lane 2, and the amplicon from cDNA in lane 3. HSP90 amplicon from gDNA is shown in lane 4, and the amplicon from the cDNA in lane 5. The first lane was loaded with 5 μL of 1 kb plus DNA ladder. All electrophoresis was performed with 10 μL of PCR products loaded per lane.

**Figure S7**

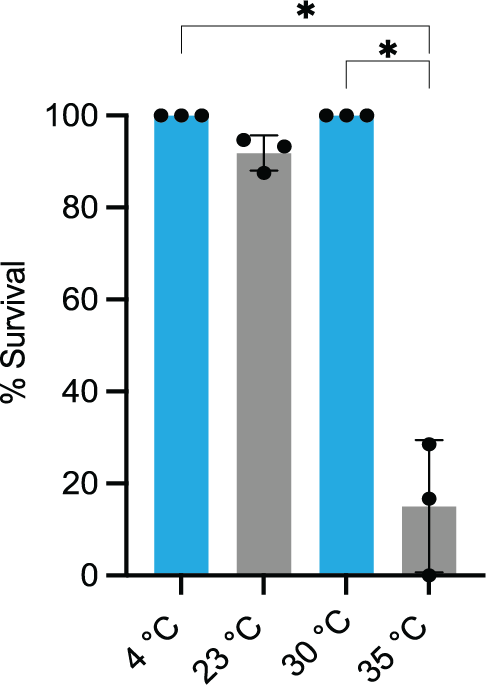

**Figure S7. Tardigrade survival rate after 24 hours of heat-stress at various temperatures.** survival rate of tardigrades treated for 24 hours at the indicated temperature. Each data point represents percent survival rate of one of three individual trials, each containing approximately 20 tardigrades. Kruskal-Wallis Test with Dunn’s multiple comparisons test, *, p= 0.0392 error bars reflect SD.

**Figure S8**

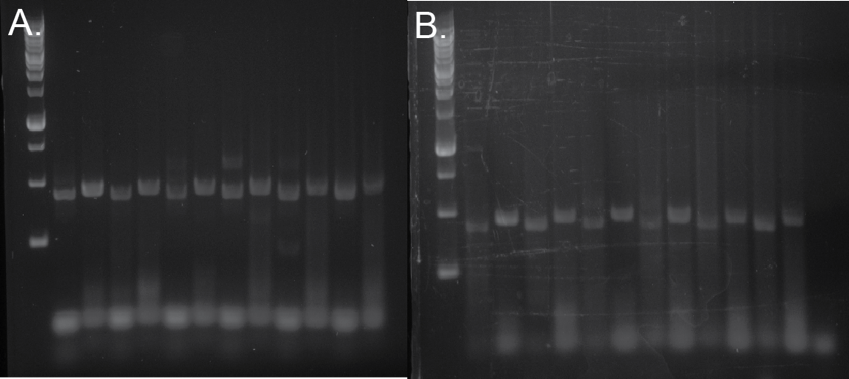

**Figure S8. Comparison of heat-shock-induced HSP70 and HSP90 transcript expression.**

a) Gel electrophoresis of HSP70 PCR products of cDNA from tardigrade samples kept at 4°C (lanes 2, 4, 6) and 30°C (lanes 8, 10, 12) for 24 hrs. For each sample, PCR products of actin were run one lane to the left of each lane containing the target gene product (lane 3, 5, 7, 9, 11, 13). b) Gel electrophoresis of HSP90 PCR products of cDNA from tardigrade samples kept at 4°C (lane 2, 4, 6) versus 30 °C (lane 8, 10, 12) for 24 hrs. Actin sample was run one lane to the left of each lane containing target gene product (lane 3, 5, 7, 9, 11, 13). The first lane was loaded with 5 uL of 1 kb plus DNA ladder. All electrophoresis was performed with 10 uL of PCR products loaded per lane.

**Figure S9**

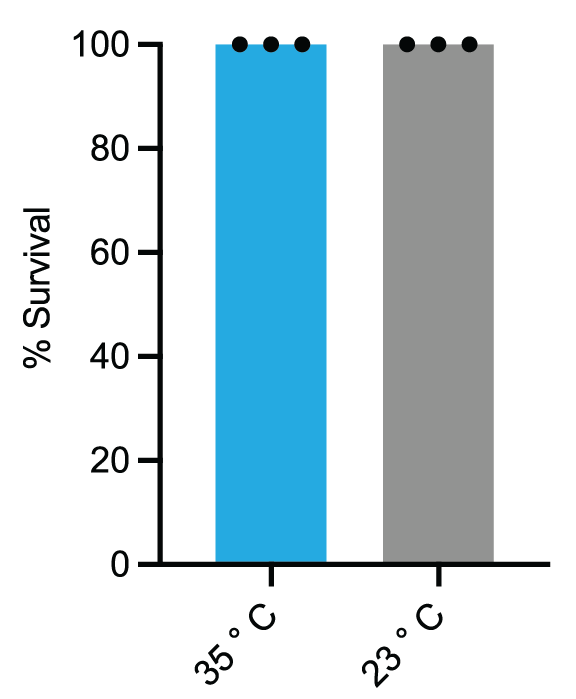

**Figure S9. Survival rates of tardigrades under heat-shock conditions.** Survival rates of tardigrades (n=15) were exposed to 20 min heat-shock at 35°C *versus* tardigrades that had been kept at room temperature (23°C) for 20 min error bars reflect SD.
